# Glucocerebrosidase Deficiency Dysregulates Human Astrocyte Lipid Metabolism

**DOI:** 10.1101/2025.01.09.632210

**Authors:** Aboud Tahanis, Thao Nguyen, Suki Oji, Marisela Martinez de Kraatz, Jazmine Jayasi, Morgan Anderson, Robert Krencik

**Affiliations:** Center for Neuroregeneration, Department of Neurosurgery, Houston Methodist Research Institute, Houston, TX, 77030, USA

**Keywords:** Parkinson’s disease, GBA1, astrocytes, human pluripotent stem cells, glycophospholipid, inflammation, lipid metabolism

## Abstract

**Background:** Deficiency in the lysosomal enzyme, glucocerebrosidase (GCase), caused by mutations in the GBA1 gene, is the most common genetic risk factor for Parkinson’s disease (PD). However, the consequence of reduced enzyme activity within neural cell sub-types remains ambiguous. Thus, the purpose of this study was to define the effect of GCase deficiency specifically in human astrocytes and test their non-cell autonomous influence upon dopaminergic neurons in a midbrain organoid model of PD.

**Methods:** Wild-type (GBA^+/+^), N370S mutant (GBA^+/N370^), and GBA1 knockout (GBA^-/-^) astrocytes were rapidly and directly induced from human pluripotent stem cells (hPSCs) via transcription factor-based differentiation. These astrocytes were extensively characterized for GCase-dependent phenotypes using immunocytochemistry, organoid coculture, enzymatic assays, lipid tracers, transcriptomics, and lipidomics.

**Results:** hPSC lines were rapidly induced into astrocytes and enzymatic assays confirmed that GBA^-/-^ astrocytes completely lacked GCase activity, while GBA^+/N370^ preserved partial activity. GBA^-/-^, but not GBA^+/N370S^, exhibited lysosomal alterations, with enlarged lysosomes and glucosylceramide (GlcCer) accumulation. GCase deficiency also exacerbated TNF-alpha-induced secretion of the inflammatory biomarker, CCL2. In midbrain organoids, GCase activity did not modulate the ability of astrocytes to support dopamine neuron production and survival. Lipidomics revealed a GBA^-/-^-specific increase in sphingomyelin, and a decrease of triglycerides. Direct rescue of GCase activity with GBA1 mRNA treatment reduced GlcCer accumulation. Astrocytes exhibited a relatively high uptake and storage of fatty acid analogs as lipid droplets, in comparison to neurons, and this process was impaired in GBA^-/-^ astrocytes. Lastly, GBA^-/-^ astrocytes accumulate neuronal membrane-derived GlcCer. These findings highlight the critical role of astrocytic GCase in lipid metabolism and its neuronal influence.

**Conclusion:** GCase deficiency does not inhibit human astrocyte differentiation nor cause a non-cell autonomous neurotoxic effect upon dopaminergic neurons within midbrain organoids. However, it does elicit enhanced inflammatory reactivity, accumulation of GlcCer, and a distinct lipidomic profile, indicating impaired lipid metabolism in astrocytes that can dysregulate neuron-astrocyte intercellular signaling. Overall, these insights underscore dysfunctional astrocyte lipid metabolism as a high priority therapeutic target in Parkinson’s disease and related neurodegenerative disorders.

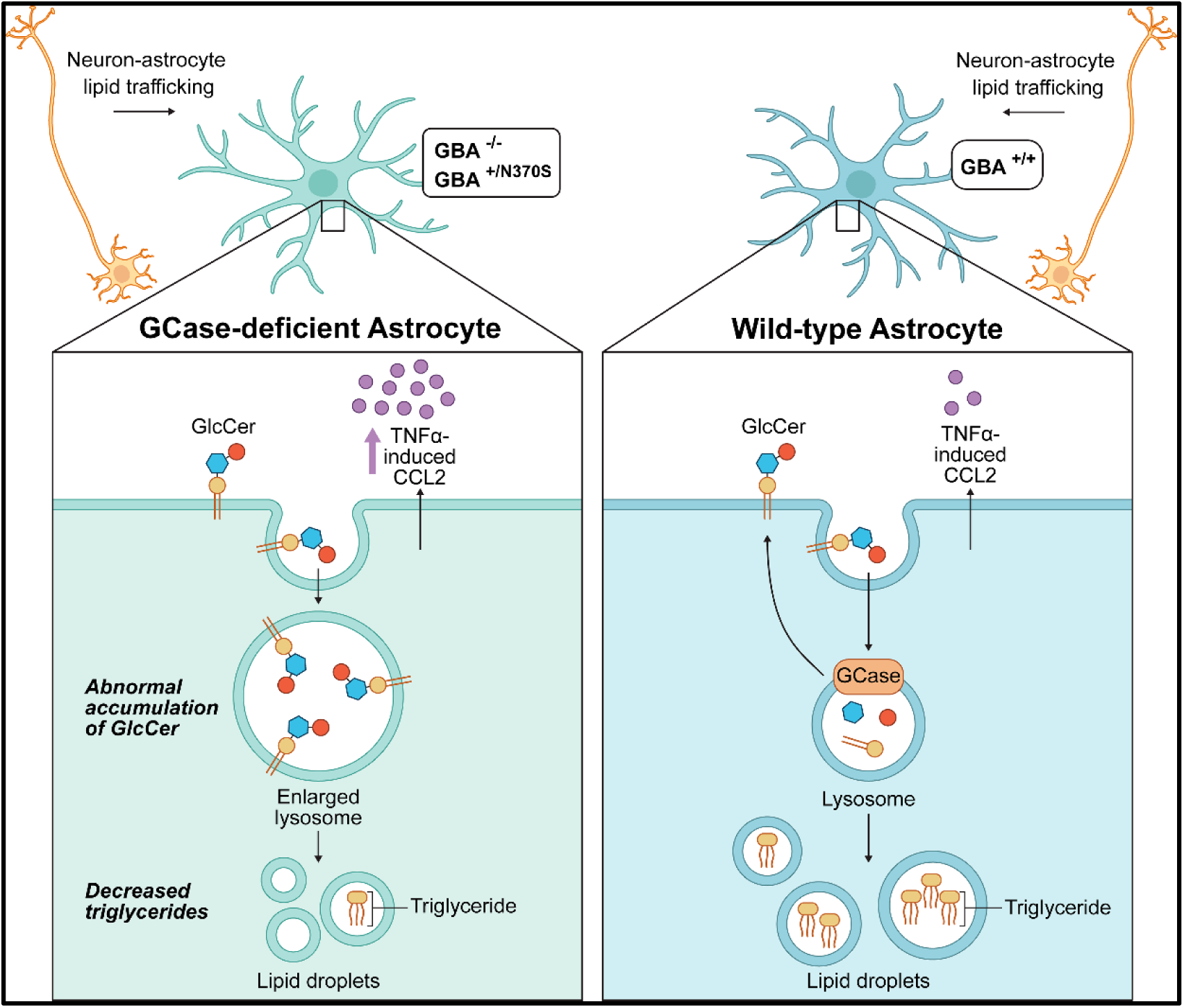

## BACKGROUND

Astrocytes are a powerhouse and recycling center in the central nervous system that contribute to neuronal activity homeostasis in the healthy brain via uptake and release of intercellular signals (e.g., synaptogenic factors, neurotransmitters, energetic substrates, antioxidants, pro/anti-inflammatories, lipids, etc.) (1–5). During neurodegeneration, astrocytes are implicated to contribute in the initiation and progression of disease through loss of supportive functions and/or gain of detrimental functions (6–8). Astrocytes, for example, protect neurons from toxic fatty acids (FAs) by uptake and metabolism during hyperactivity (9) and oxidative stress (10). On the other hand, disease-associated reactive astrocytes can abnormally accumulate (11, 12) or secrete (13) FAs that may lead to altered neuronal activity or reduced viability. However, a comprehensive understanding of key membrane components and the extent of intercellular dependency for human astrocyte lipid metabolism remains lacking in healthy and diseased states.

Parkinson’s disease (PD) is a progressive neurodegenerative disorder characterized by the loss of dopaminergic (DA) neurons in the substantia nigra pars compacta in the midbrain. This results in the trilogy of motor symptoms: bradykinesia, rigidity, and tremor. Both genetic and environmental factors are thought to contribute to the development of the disease (14). Research into the underlying cellular consequence of genetic mutations has identified converging dysfunctional pathways in astrocytes including inflammation, organelle health (i.e., mitochondria and lysosomes) and lipid handling (15–17). Given that lipid metabolism dysfunction is observed in human PD brains (18–20), a better understanding of human astrocyte lipid metabolism is needed to uncover potential underlying causes and therapeutic targets for PD. Mutations in the GBA1 gene (encoding a glycolipid metabolism enzyme in the lysosome, glucocerebrosidase [GCase]) are the most common monogenic risk factors for PD and are associated with both dementia with Lewy bodies and frontotemporal dementia. Biallelic mutations result in the lysosome storage disorder Gaucher disease with or without Parkinsonian features (21). Research of patient tissue and experimental models have revealed that reduced GCase activity leads to accumulation of the lipid substrate glucosylceramide (GlcCer), and this has been linked to multiple downstream cellular phenotypes including organelle dysfunction and increased α-synuclein aggregation (22, 23). Specifically related to astrocytes, immature human astrocytes generated from human pluripotent stem cells (hPSCs) were reported to exhibit multiple phenotypes including increased GlcCer and glucosylsphingosine content, and increased secretion of a subset of inflammatory molecules in baseline conditions (24). However, an extensive profiling of lipids and whether GCase-deficiency in mature astrocytes causes a non-cell autonomous neurotoxic degeneration upon human DA neurons has yet to be examined.

We previously reported a protocol for rapid differentiation of mature astrocytes from hPSCs using inducible transcription factors SRY-box 9 (*Sox9*) and nuclear factor 1 A (*NFIA*), and a protocol for incorporation of astrocytes into neural organoids as a well-defined and reproducible experimental platform (25). Here, we expand upon this approach to rapidly generate human astrocytes from hPSCs with GBA1 mutations for comprehensive GCase-dependent phenotyping and to incorporate them into midbrain organoids for assessing the consequence upon DA neurons. Our results conclude that GCase deficiency does not interfere with astrocyte differentiation and does not generate overtly neurotoxic astrocytes. However, these astrocytes do exhibit intracellular accumulation of neuron-derived GlcCer, enhanced inflammatory reactivity, and a dysregulated lipidomic, altogether providing novel insights into cellular dysfunction due to GCase deficiency.

## RESULTS

### Glucocerebrosidase-deficient human astrocytes can be rapidly generated from hPSCs

First, we analyzed RNA sequencing data we previously generated from hPSC-induced neural progenitor cells (NPCs), astrocytes, and neurons (25) to identify PD-linked candidates that are highly and relatively abundant in astrocytes. PARK7 (encoding DJ-1 protein) was the most abundant transcript among all three cellular subtypes. On the other hand, alpha-synuclein (SNCA), which regulates synaptic vesicle clustering, docking, and the release of neurotransmitters from the presynaptic terminal, was overwhelmingly expressed in neurons. GBA1 transcripts were abundant and relatively restricted to astrocytes (**Fig. 1A**). Because GBA1 mutations are the highest risk factor of PD, we sought to subsequently analyze their effect in astrocytes to determine how it might contribute to the progression of PD. Thus, we genetically engineered two biologically distinct mutant GBA1 (GBA^+/N370S^) hPSC lines, derived from PD patients, with an inducible Sox9-NFIA transgene for rapid production of astrocytes as previously described (25). Further, using a previously characterized wild-type line containing the Sox9-NFIA transgene, we generated a Cas9-mediated complete isogenic knockout (KO) line (GBA^-/-^) that was verified by sequencing and immunocytochemistry (ICC) (**Fig. 1B, Fig. S1A, B**). To determine activity levels of the GCase enzyme, we utilized a cell-permeable substrate 5-(pentafluorobenzoylamino)fluorescein Di-β-D-glucopyranoside (PFB-FDGlu) that fluoresces upon being metabolized by GCase. As a positive control for GCase deficiency, cells were treated with a GCase irreversible inhibitor, conduritol-β-epoxide (CBE). No activity was detected in GBA^-/-^ astrocytes nor in those treated with CBE, whereas GBA^+/N370S^ exhibited 33.9% (SD=6.01%) of wildtype activity (**Fig. 1C**). Identity of astrocytes at 15 days of induction was confirmed by detection of astrocyte markers CD44 and GFAP (**Fig. 1D, Fig. S1C).** Stellate astrocyte-like morphologies were observed with high variability in size, branching patterns, and staining intensity within and among all groups as expected. Finally, because GCase is primarily active within lysosomes, we analyzed astrocyte lysosomal content with Lysotracker-based live imaging. This revealed significantly enlarged lysosomes in GBA^-/-^ astrocytes, but not GBA^+/N370S^, in comparison to GBA^+/+^. These findings indicate GCase deficiency does not interfere with astrocyte differentiation, yet complete KO leads to disruption of lysosomal homeostasis, potentially through the accumulation of lipid substrates.

**Figure 1.**
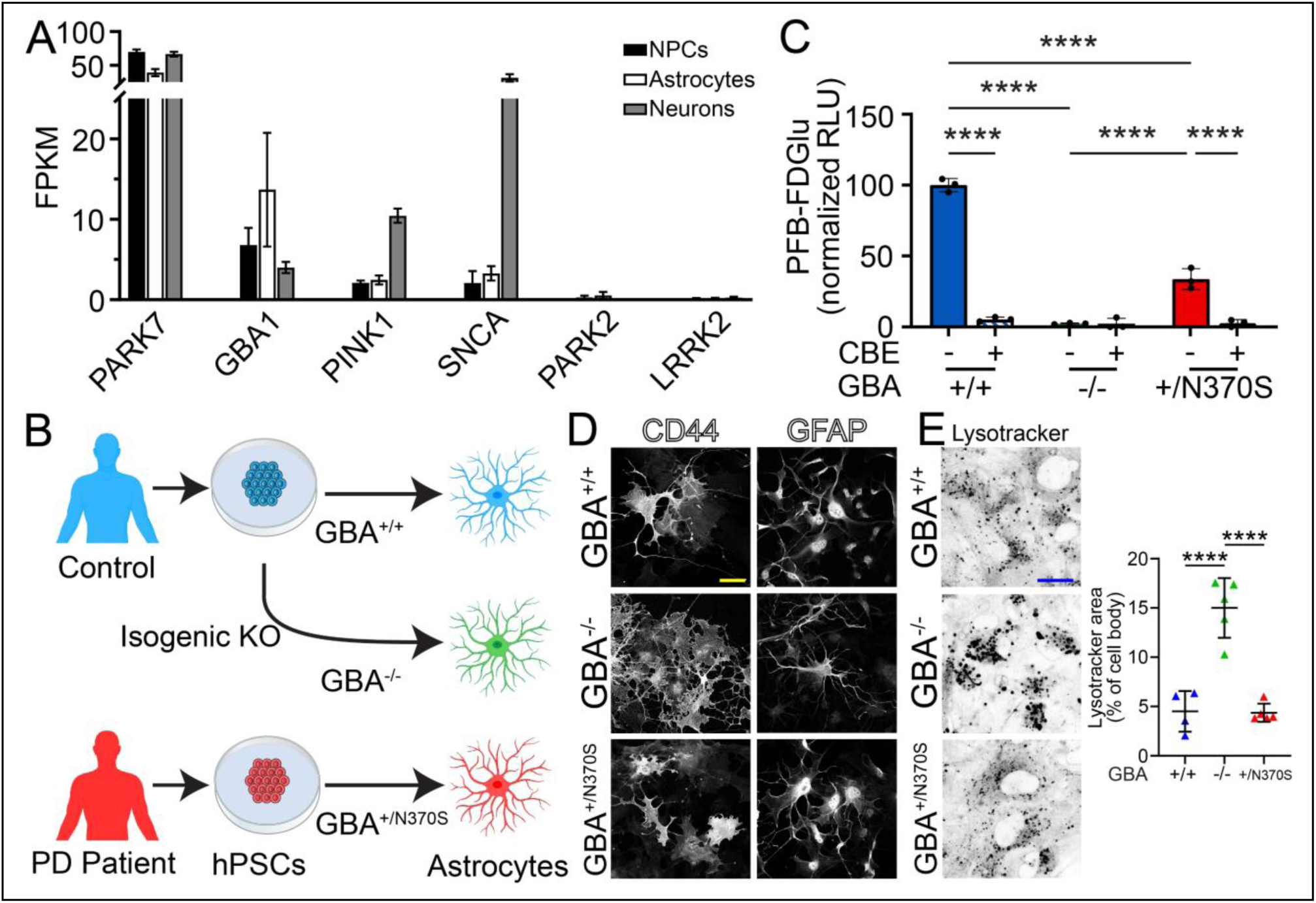
Glucocerebrosidase-deficient human astrocytes can be rapidly generated from hPSCs. (A) RNA sequencing reveals relatively high expression of GBA1 in hPSC-astrocyte. (B) A schematic diagram depicting the workflow to derive cells from healthy individuals, GBA1 knockouts, and PD patients. (C) Quantification of the GCase activity levels based on the fluorescent signal of PGB-FGDGlu with and without treatment with conduritol-β-epoxide (CBE) treatment, an irreversible inhibitor of GCase. (D) Fluorescent images verifying expression of astrocytes markers CD44 and GFAP, following the induced differentiation process. (E) Representative images of live cell staining with Lysotracker showing enlarged lysosomes in GBA^-/-^ compared to GBA^+/+^ (left). Scale bar: 50µm. Quantification of lysosome size (right). Scale bar: 20µm. **** represents p<.0001 and data points represent replicate studies.

### Astrocytes support dopaminergic neurons in midbrain organoids independent of glucocerebrosidase activity

Reactive astrocytes can cause non-cell autonomous degeneration upon neurons, including DA neurons, either through a loss of neuroprotection or gain of neurotoxic substances through a variety of different mechanisms (26). To investigate this phenomenon in GCase-deficient human astrocytes, we first assessed their reactivity to the inflammatory cytokine TNFα, using secreted CCL2 as a biomarker as previously described (25). 48 hours after 10ng/ml treatment, astrocytes increased secretion of CCL2 regardless of the genotype, with a significant increase in both GBA^+/N370S^ and GBA^-/-^ astrocytes compared to GBA^+/+^ **(Fig. 2A)**. To test the potential consequence of this GCase-dependent increased inflammation upon human DA neurons, we first optimized differentiation of midbrain organoids by comparing different supplement conditions previously utilized to generate hPSC-derived midbrain DA neurons (27), and by testing whether combinatorial induced expression of transcription factors ASCL1, LMX1B, and NURR1 (together, ALN) can enhance DA neuron production (28) **(Fig. 2B)**. hPSCs were differentiated for 20 days as monolayer cultures, then combined into free-floated organoids at approximately 1,700 cells per organoid using method previously detailed (29). At day 40 of differentiation, the highest yield of DA neurons, determined by counting TH+ neurons per organoid with ICC, was achieved by combinatorial addition of TGFβ and dbcAMP **(Fig. 2C, D, Fig. S2A)**, representing ∼15% of the total population. Interestingly, cell bodies reproducibly cluster at one end of the organoid while fibers extensively project throughout the rest of the organoid, indicating self -organization over time. Even though doxycycline was able to induce expression of the ALN transgenes as well as downstream midbrain progenitor transcripts FoxA1 and Lmx1 (**Fig. S2B**), and transgene induction led to production of a small amount of TH+ neurons in monolayer cultures (not shown), this strategy did not significantly increase the number of TH+ neurons within midbrain organoids. Next, we tested whether GCase-deficient inflammatory astrocytes displayed a loss of support, or neurotoxic effect, while cocultured as 20% of the total cellular population of midbrain organoids starting on day 20. At day 40 endpoint, we confirmed a homogenous distribution of CD44+ astrocytes throughout the midbrain organoids. ICC revealed significantly higher number of TH+ DA neurons when cocultured with astrocytes, indicating a supportive role, and no significant difference between organoids in the presence of GBA^+/+^, GBA^+/N370S^ and GBA^-/-^ astrocytes, with and without chronic 14 days of treatment with TNFα. Despite the increased inflammatory response in GCase-deficient astrocytes, no toxic effect on the total number of DA neurons in optimal conditions was observed **(Fig. 2E, F)**.

**Figure 2.**
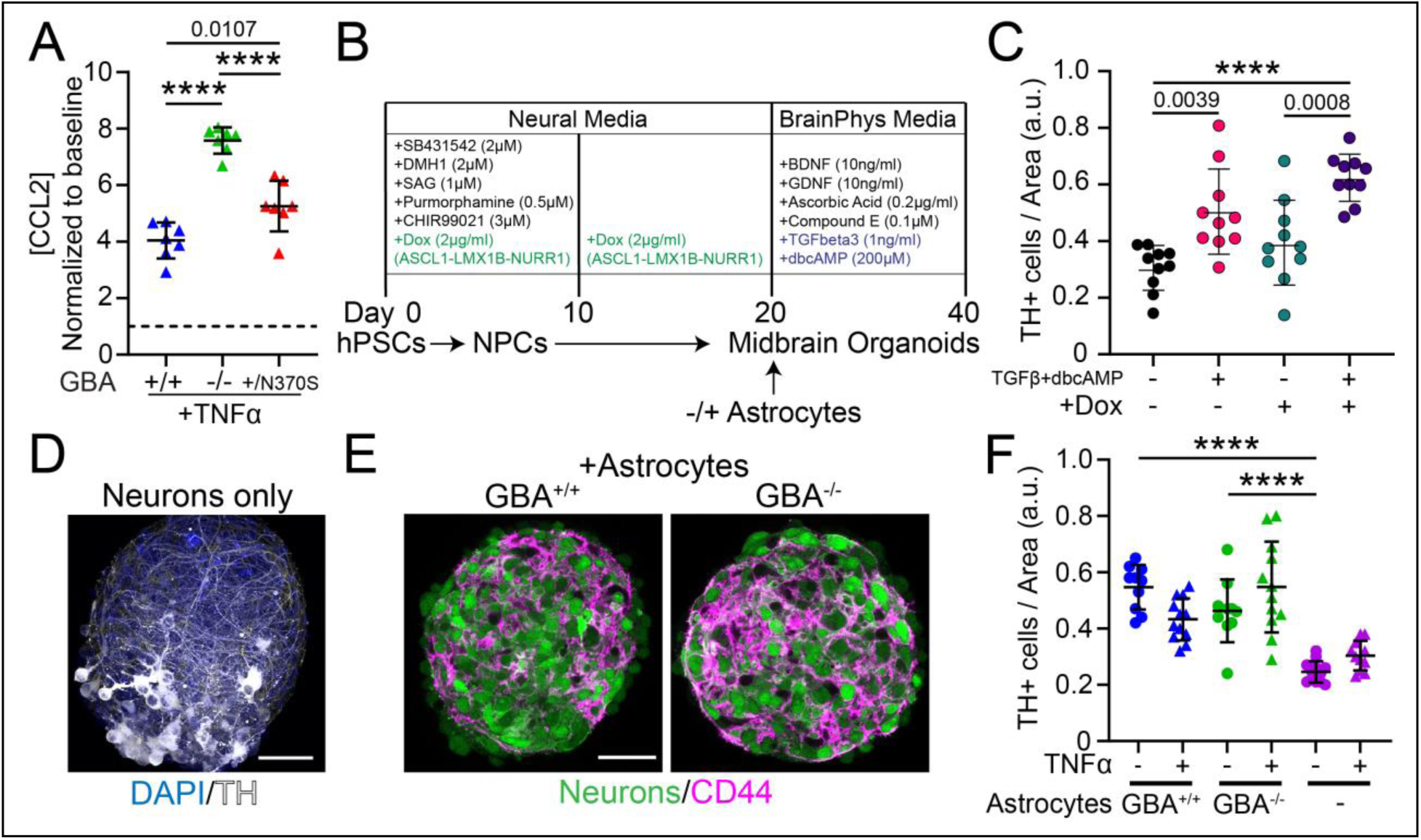
Astrocytes improve the yield of dopaminergic neurons in midbrain organoids and do not exhibit neurotoxicity. (A) Secretion of CCL2 by GBA^+/+^, GBA^+/N370S^, and GBA^-/-^ astrocytes after treatment with TNF-alpha. (B) Timelines of midbrain organoid differentiation protocols tested for the optimization of dopaminergic (DA) neuron differentiation. (C) Counts of tyrosine hydroxylase (TH)-positive DA neurons comparing different differentiation protocols. (D) Representative ICC image of differentiated DA neurons in midbrain organoids. (E) Representative image of midbrain organoids composed of neurons (GFP) and astrocytes (CD44). (F) DA neuron counts in midbrain organoids with and without astrocytes under inflammatory conditions.

### Glucocerebrosidase deficiency elicits a reactive astrocyte lipidome

To investigate astrocyte baseline reactivity in the context of GCase deficiency, we performed RNA sequencing analysis of GBA^-/-^ astrocytes and compared to GBA^+/+^ under inflammatory conditions. The transcriptomic level of key inflammatory biomarkers of hPSC-astrocytes, including CXCL8, LIF, CCL2, CTGF, and THBS1 (25), were not upregulated at baseline conditions in GBA^-/-^ astrocytes. However, GFAP was upregulated, along with several extracellular matrix-related components such as ACAN, suggesting there is a baseline fibrotic-like reactivity state **(Fig. 3A, Fig. S3A)**. When treated for two days with interleukin-1a, TNFα, and complement C1q (a.k.a., ITC) similar as previously described to generate astrocytes with a reactive transcriptomic and lipidomic signature (13), GBA^-/-^ astrocytes did significantly upregulate numerous inflammatory transcripts in respond to the exogenous inflammatory cytokines **(Fig. S3B)**. Next, we conducted unbiased lipidomic profiling including 920 lipid species across 14 different classes (**Sup. Data Table 1**). We specified neural progenitors to the midbrain lineage with a combination of Sonic hedgehog and a Wnt agonist (27) prior to doxycycline-induced Sox9-NFIA induction, producing En1-positive astrocytes (**Fig. S3C**). Lipidomics revealed no significant differences in the total concentrations of most lipid classes including phosphatidylcholine (PC), phosphatidylethanolamine (PE), phosphatidylserine (PS), phosphatidic acid (PA), and phosphatidylglycerol (PG). However, there were significant differences in the sphingomyelin (SM) and triglyceride (TG) classes. Specifically, we observed an increase in SM levels in GBA^-/-^ astrocytes compared to GBA^+/+^ and GBA^+/N370S^. SM, similar to glycophospholipids, are membrane components and that traffic to the lysosomes to be broken down into ceramides. High SM content in GBA^-/-^ cells indicates a disrupted lysosomal function beyond GCase as was previously observed in post-mortem brain tissue of PD patients (30) . Conversely, the total TG content, which represents the storage form of FA as lipid particles, was decreased in GBA^-/-^ astrocytes **(Fig. 3B)**. Interestingly, a similar pattern of decreased TG was observed in GBA^+/+^ astrocytes following activation by inflammatory cytokines **(Fig. 3C)**, while SM levels were not altered by inflammation. Principal component analysis determined that GBA^-/-^ astrocytes have a distinct lipid cluster from that of GBA^+/+^ and that this pattern did not change with treatment of inflammatory mediators **(Fig. 3D)**. GBA^+/N370S^ astrocytes, on the other hand, had a similar lipidomic profile to GBA^+/+^ **(Fig. S3D)**.

**Figure 3.**
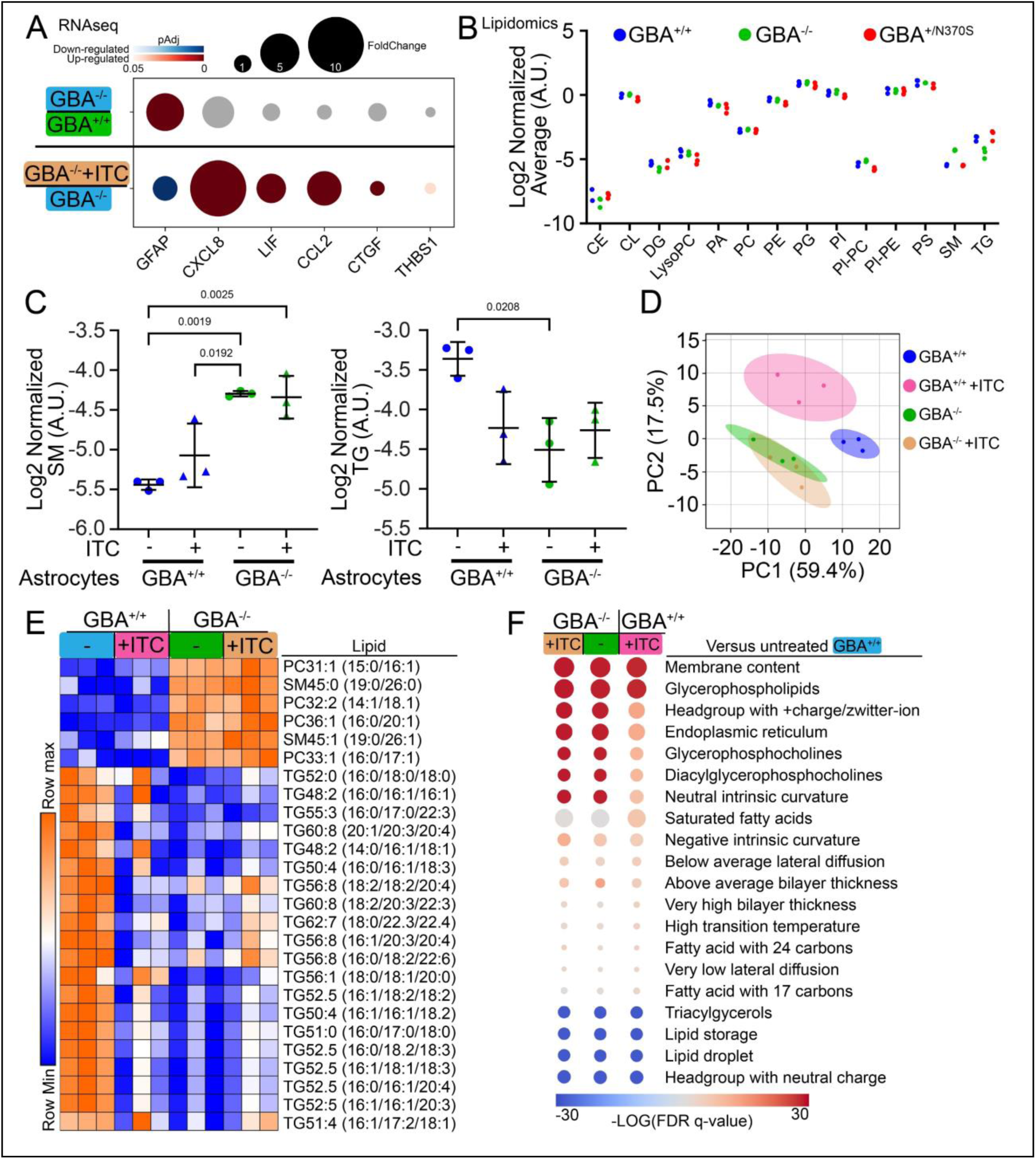
Glucocerebrosidase deficiency elicits a reactive astrocyte lipidome. (A) Transcriptomic analysis of key inflammatory genes in GBA^-/-^ vs GBA^+/+^ astrocytes at baseline, and with inflammatory cytokines treatment. GBA^-/-^ upregulated inflammatory genes after treatment. (B) Lipid class analysis indicates no difference in major phospholipid classes but shows a significant increase in sphingomyelins (SM) and a decrease in triglycerides (TG) in GBA^-/-^ astrocytes. (C) SM and TG levels in astrocytes at baseline and in inflammatory conditions reveal increased SM in GBA^-/-^ and reduced TG, similar to the levels seen in GBA^-/-^ Astrocytes (D) Principal component analysis of lipidomic profiles demonstrates a distinct lipid composition in GBA^-/-^ astrocytes compared to GBA^+/+^. (E) Heatmap of the top differentially abundant lipids between GBA^-/-^ and GBA^+/+^ astrocytes. (F) Lipid enrichment analysis of GBA^-/-^ and inflamed GBA^+/+^ compared to baseline.

Comparison of individual lipids that exhibit the highest difference between groups (i.e., greater than 1.5-fold and significantly different) revealed lipids that are increased in GBA^-/-^ (at both baseline and inflammatory conditions) in comparison to GBA^+/+^ (**Fig. 3E**). Interestingly, these same lipids are not changed in inflammatory GBA^+/+^, though they do exhibit a decrease in some individual TGs. Finally, to go beyond comparing concentrations of lipids species and lipid classes, we applied lipid ontology analysis to classify and compare astrocytes based on lipid function (31). In comparison to untreated GBA^+/+^ astrocytes, GBA^-/-^ exhibit an increase in membrane components and glycophospholipids, coupled with decrease in triacylglycerols and associated lipid droplet, among others, both with and without cytokine treatment (**Fig. 3F**). In conclusion, we identified a GCase-dependent lipidome that is not evident by solely observing transcriptional profiles and we discovered both distinct and overlapping with features with inflammatory astrocytes.

### Glucocerebrosidase-deficient astrocytes accumulate GlcCer that can be restored with GBA1 mRNA

To specifically investigate the handling of glycophospholipids, we treated astrocytes with fluorescent-labeled GlcCer (NBD-GlcCer) for 30 min, followed by a complete media exchange. Similar to native GlcCer, exogenously NBD-GlcCer integrates into the cellular membrane, then recycles to the lysosome via endocytosis where it is catalyzed by GCase, resulting in the loss of fluorescent signal. 24-hour after treatment, NBD-GlcCer significantly accumulated within GBA^-/-^ astrocytes in comparison to GBA^+/+^, but not in GBA^+/N370S^ (**Fig. 4A**). This indicates that complete GCase deficiency overtly disrupts lipid metabolism while low amounts of active GCase activity in GBA^+/N370S^ is still sufficient to avoid this phenomenon in astrocytes. We next aimed to increase GCase activity and rescue the lipid accumulation. First, we tested astrocyte treatment with ambroxol (ABX), a small molecule mucolytic that has been reported to increase GCase activity in the brain in preclinical experiments (32, 33) and is currently in clinical trials for PD (34). We assessed whether ABX (10µM, based on prior studies) for 48 hours reduced accumulation of NBD-GlcCer in GBA^-/-^ astrocytes. No difference was observed, potentially because ABX may exert its action by binding to GCase protein (35) (**Fig. 4B**). Thus, we tested whether ABX enhances GCase activity in GBA^+/N370S^ astrocytes. Extracellular lactate dehydrogenase (LDH) levels were used to determine tolerable ABX concentrations (**Fig. 4C**). Surprisingly, GCase activity was inhibited by ABX, indicating that while ABX may have potential in increasing GCase activity in other cell types, its application in astrocytes appears limited, possibly because of technical differences in culture conditions or cell sub-type dose tolerance (**Fig. 4C, D)**. As an alternative approach to restore GCase activity, and to directly confirm whether the NBD-GlcCer accumulation in GBA^-/-^ astrocytes is specific to the loss of activity, we tested the potential to rescue with GBA1 mRNA transfection (**Fig 4E**). Remarkably, NBD-GlcCer accumulation in GBA^-/-^ astrocytes was significantly decreased 24 hours after mRNA treatment, partially restoring clearance towards levels observed in GBA^+/+^ (**Fig. 4F**). Together, these results highlight the role of GCase in maintaining lipid homeostasis and proper metabolism in astrocytes, as its deficiency disrupts the recycling of plasma membrane components leading to the accumulation of GlcCer and altering the overall lipid composition.

**Figure 4.**
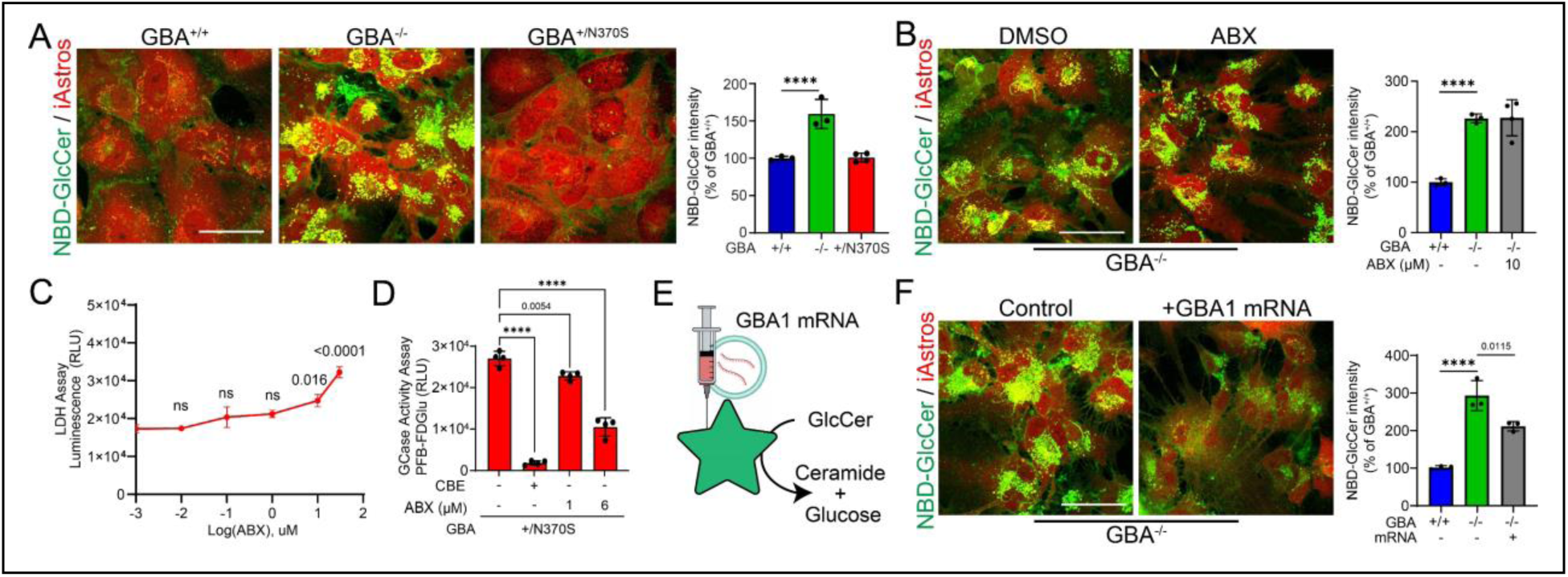
Glucocerebrosidase-deficient astrocytes accumulate GlcCer. (A) NBD-GlcCer significantly accumulates within GBA^-/-^ deficient astrocytes. (B) Ambroxol (ABX) treatment does not modulate NBD-GlcCer accumulation. (C) Viability is reduced by ABX treatment at 10µM and above as indicated by increased LDH secretion. (D) GCase activity of mutant astrocytes does not increase in response to ABX treatment. (E) Schematic of GBA1 mRNA transfection. (F) GBA1 mRNA transfection can rescue NBD-GlcCer accumulation in GBA-/- cells. Data points represent replicate studies. **** represents p<.0001.

### Neuron-to-astrocyte lipid metabolism is disrupted by glucocerebrosidase deficiency in astrocytes

To determine how astrocyte lipid handling is altered by neurons and whether astrocyte GCase deficiency alters neuron-lipid metabolism (**Fig. 5A**), we measured the uptake of FAs by astrocytes and the intercellular transfer of GlcCer between neurons and astrocytes. We first treated GBA^+/+^ astrocytes and neurons with 5μM of C1-BODIPY-C12, a fluorescent FA analog for 48h. FAs can be taken from the external environment and stored as TG within cytoplasmic compartments, called lipid droplets, for energy stores and other purposes (36). Results indicated that FA uptake and/or storage uptake is relatively much higher in astrocytes in comparison to neurons in both midbrain organoids and monocultures **(Fig. 5B, C)**. Furthermore, astrocytes exhibited an increase in lipid droplets size and number in the presence of neurons in monolayer cocultures. To test whether this is influenced by GCase deficiency, we treated astrocyte monocultures with C1-BODIPY-C12. In accordance with lipidomics data, GBA^-/-^ astrocytes in coculture with neurons displayed a significant reduction in lipid storage as compared to GBA^+/+^ and GBA^+/N370S^ astrocytes **(Fig. 5D)**. Finally, to investigate whether GCase deficiency leads to dysregulated metabolism of lipids that originate directly from neuronal membranes, similar as previously described in *Drosophila* (37), we first directly treated neurons with NBD-GlcCer for 30min. Subsequently, the media was replaced, and astrocytes were directly cocultured with neurons for an additional 24 hours. This revealed that GBA^-/-^ astrocytes significantly accumulated neuron-derived NBD-GlcCer in comparison to GBA^+/+^ and GBA^+/N370S^, indicating a dysregulated neuronal lipid metabolism.

**Figure 5.**
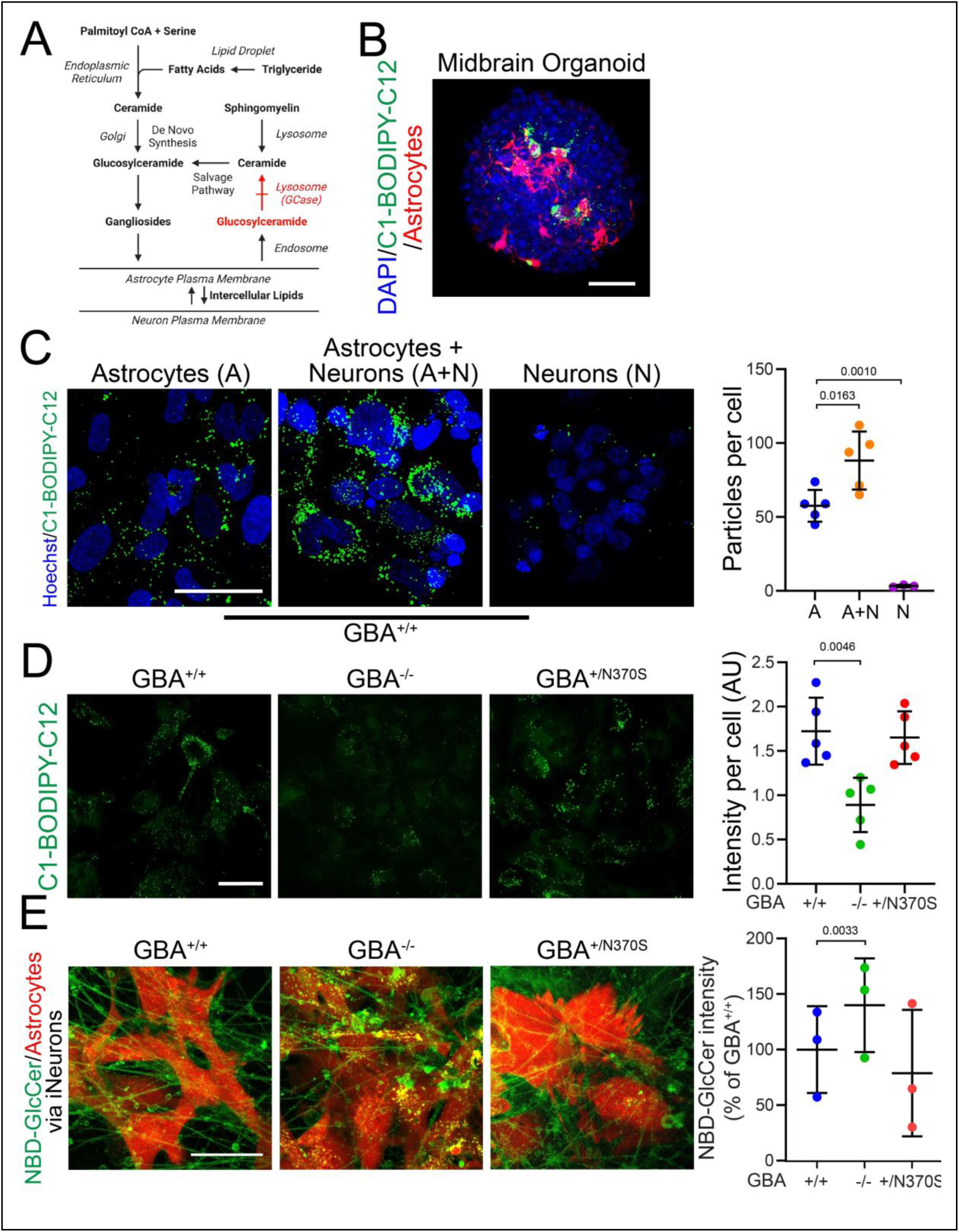
The presence of neurons and neuron-derived GlcCer reveals dysregulation in GCase-deficient astrocyte lipid storage and metabolism. (A) Schematic of neuron lipid storage and metabolism in astrocytes. (B) C1-BODIPY-C12 uptake in 3D organoids demonstrates fatty acid storage in astrocytes. (C) Representative images of C1-BODIPY-C12 uptake in astrocytes, neurons, and co-cultures of both cell types reveal that astrocytes are the primary cells responsible for fatty acid storage. Uptake is increased in astrocytes co-cultured with neurons, while neurons exhibit minimal storage of fatty acids (left). Quantification of fluorescent lipid droplets per cell (right). (D) C1-BODIPY-C12 uptake in astrocytes exhibits a significant reduction in uptake in GBA^-/-^ astrocytes compared to GBA^+/+^ (left). Quantification of fluorescence intensity per cell (right). (E) NBD-GlcCer trafficking from neurons to astrocytes shows lysosomal accumulation in GBA^-/-^ astrocytes(left). Quantification of NBD-GlcCer fluorescence intensity in astrocytes (right).

## DISCUSSION

Astrocyte cellular membranes, lipid metabolism, and lipid rafts are increasing becoming recognized as important mediators in autonomous function and non-cell autonomous signaling during neurodegeneration (38, 39) with potential toxic (13) or protective effects (9, 40). In this study, we uncovered that at least partial GCase activity is essential to maintain proper lipid composition in cultured human astrocytes. This was confirmed through observing rapid rescue of GlcCer accumulation via GBA1 mRNA delivery to elicit GCase activity in GBA^-/-^ astrocytes. Though partial GCase-deficiency in mutant astrocytes did not display overt lipid dysregulation, they did exhibit an enhanced inflammatory reactivity, suggesting that the inflammatory microenvironment in the aged, diseased, and/or injured brain may exacerbate phenotypes influenced by astrocytes. This is reminiscent of mouse models of GCase deficiency that exhibit systemic inflammation outside of the nervous system in the absence of major GlcCer accumulation (41). Clinical symptoms may also vary depending on cell subtype-specific responses. For example, within the nervous system, microglia-specific KO mice lack GlcCer accumulation and disease symptoms, yet neuron-specific KO do exhibit these phenotypes as well as gliosis (42). Thus, we expect high priority subsequent studies using the midbrain organoid approach described here should include testing whether GCase deficiency within neurons can induce astrocyte reactivity, which together with GCase deficiency in astrocytes may cause feedback loops to accelerate neurodegeneration.

To test the non-cell autonomous effect of GCase-deficient astrocytes upon dopaminergic neurons, we utilized a midbrain organoid approach that enabled us to directly incorporate post-mitotic astrocytes with neurons, while avoiding the influence of non-neural cells that typically are present in primary cell cultures or hPSC-based cultures that are differentiated over extensive time periods. Based on previous studies in PD and other neurodegenerative disorders, we expected that disease-associated astrocytes in baseline or inflammatory conditions can lead to direct neurodegeneration via secretion of molecules such as complement components or inflammatory mediators (26). However, many of these studies that investigated neurotoxicity have not directly demonstrated a toxic effect directly upon neurons and instead solely profile the inflammatory signature of astrocytes. Here, we did not detect GCase-dependent or inflammation-dependent toxicity, as defined by measuring the number of DA neurons within organoids. This lack of obvious degeneration or may be due to a variety of factors, such as the limited timeline used in this study from the start of exposure to endpoint readout, the presence of numerous pro-survival cell culture components in the media such as antioxidants that may interfere with potential toxicity, the lack of neuronal receptors in this study that can detect astrocyte-derived components, the lack of immune cells, and/or the absence of additional degeneration-inducing factors such as alpha-synuclein or oxidative stress. Regardless, this study did identify a beneficial effect of astrocytes upon DA neuron content within midbrain organoids, and the mechanisms underlying this effect may be elucidated in future studies.

GBA1 mutations are strongly implicated in both familial and sporadic PD. Post-mortem studies in patients with other genetic causes of PD, such as those carrying LRRK2 mutations, demonstrate a decrease in GCase activity even in the absence of GBA1 mutations (43, 44). Moreover, GCase activity declines with age, accompanied by increased neuroinflammation (45), underscoring the critical role of lipid metabolism in cellular homeostasis and its contribution to the proteomic hallmarks of PD (46). A key pathological feature of PD is the aggregation of alpha-synuclein, a protein that interacts with the cytoplasmic membrane bilayer (47, 48). Studies indicate that fatty acids influence both the accumulation and clearance of alpha-synuclein aggregates. However, the astrocytes generated in this study express limited or undetectable levels of alpha-synuclein, which precludes direct investigation into the effects of lipid alterations on protein misfolding without further coculture studies. Notably, our lipidomic analysis revealed an increase in sphingomyelin and related lipids, consistent with changes observed in post-mortem PD brains (30, 49). Sphingomyelin plays a pivotal role in organizing cellular membrane, modulating lipid-protein and protein-protein interactions, and regulating molecular trafficking across membranes (50). Under inflammatory conditions, these mechanisms are critical for cellular survival and the restoration of homeostasis.

Altogether, our findings demonstrate that the lack of glucocerebrosidase activity in human astrocytes causes significant disruption in lysosomal homeostasis and membrane lipid composition and function potentially contributing to the inflammation in Parkinson’s disease and other diseases. This research highlights the critical role of GBA1 mutations in disrupting lipid metabolism and inflammatory responses in human astrocytes. We demonstrated that GCase deficiency leads to abnormal accumulation of glucosylceramide and a distinct lipidomic profile, indicating impaired lipid handling and altered cellular function. Looking toward potential future clinical implications, our studies indicate that GBA1 mRNA therapy may be promising in restoring GCase activity and mitigating lipid accumulation. Overall, these insights underscore the importance of targeting astrocyte lipid metabolism as a potential therapeutic strategy in Parkinson’s disease and related neurodegenerative disorders.

## MATERIALS AND METHODS

### Cell culture

hPSC line WTC11 was obtained from Coriell Institute (#GM25256) to generate GBA^+/+^ and GBA^-/-^ astrocytes. GBA^-/-^ hPSCs were generated using sgRNAs (GCCCGUGUGAUUAGCCUGGA, GCUAUGAGAGUACACGCAGUGAGCUGUAGCCGAAGCUUUU) with Synthego Gene Knockout Kit v2 and verified with sequencing (PCR primers: forward, CCTTCACTTTCTGGAACTTCTGT, reverse, GAAACCGTGTTCAGTCTCTCCT. Sequencing primer: CCTTCACTTTCTGGAACTTCTGTT). This study also utilized PD patient-derived GBA^+/N370^ hPSCs (PPMI.I.1102.2 (RRID:CVCL_D5U0, a.k.a. 507-GBA) (PPMI.I.1084.1 (RRID:CVCL_D5SN), a.k.a 569-GBA^+/N370S^), obtained from the Parkinson’s Progression Markers Initiative (PPMI) database (https://www.ppmi-info.org/access-data-specimens/download-data), RRID:SCR_006431. For up-to-date information on the study, visit http://www.ppmi-info.org. 507-GBA^+/N370S^ astrocytes were used for all experiments, whereas 569-GBA^+/N370S^ astrocytes were used in select experiments to confirm reproducibility among distinct lines. hPSCs were maintained in Matrigel-coated 6-well plates (Fisher Scientific #353046) in TeSR-E8 basal medium with supplements (StemCell Technologies #05990) and passaged weekly using ReLeSR (StemCell Technologies #05872). To genetically engineer hPSCs, rapidly induce the differentiation of astrocytes and midbrain neurons, and culture over time, we utilized our previously detailed TALE nuclease and culture protocol (25).

### NBD-GlcCer and C1-BODIPY-C12 cell labelling

Astrocytes were plated onto imaging slides (Ibidi #80826), precoated with Matrigel, at a density of 40,000 cells per well. To test the accumulation of glycophospholipid, the cells were incubated with 4µM C6 NBD-GlcCer (Cayman Chemical #23209) for 30 minutes to allow insertion in the plasma membrane. The cells were then incubated in NBD-GlcCer-free media for 18-24 hours to allow for internalization and fluorescent confocal imaging (Leica DMi8). For cocultures studies, NBD-GlcCer colocalized with the astrocyte mApple reporter was specifically measured within confocal image slices to avoid signal from neuronal membrane. To investigate the storage of TG, cells were incubated for 48 hours with 5μM of fluorescent fatty acid analog C1-BODIPY 500/510 C12 (a.k.a., C1-BODIPY-C12, ThermoFisher #D3823). The chemical was suspended in ethanol at 5mM and stored in -20 until use.

### mRNA transfection

GBA1 mRNA was produced by the RNAcore at Houston Methodist Research Institute and reconstituted to a concentration of 1 μg/μl in water. GBA1 mRNA or a water control was delivered to the cells using the JetMessenger transfection kit (Polyplus #101000056), following the manufacturer’s instructions. 12 hours post-transfection, the cells were treated with NBD-GluCer as previously described and imaged after an additional 12 hours.

### Viability and toxicity assays

To study the toxicity of ambroxol (Millipore-Sigma #PHR2063), LDH-Glo Cytotoxicity Assay (Promega, #J2380) was performed. Astrocytes were treated with varying concentration for 48 hours. 5 µl of cell culture media was collected, transferred into a white 96-well plate and diluted 10 times with 45 µl of PBS. 50µl of LDH Detection Reagent, prepared as described by the manufacturer, was added to each sample in the 96-well plate, and the plate was incubated for 1 hour at room temperature. The luminescence signal was the measured using Tecan microwell reader.

### Glucocerebrosidase activity assay

Cells were plated in a 96-well plate at a density of 25,000 cells per well with four technical replicates and allowed to attach for 2 hours. Following attachment, the cells were re-fed with phenol red-free medium, with or without the addition of 4 μM CBE (Sigma, #C5424) for 48 hours. PFB-FDGlu (5-(Pentafluorobenzoylamino)Fluorescein Di-β-D-Glucopyranoside) (ThermoFisher #P11947) substrate was added at a final concentration of 375 μM. The cells were further incubated for 1 hour to allow the uptake of PFB-FDGlu via pinocytosis and its subsequent accumulation into the lysosome. The fluorescent signal was detected using a microplate reader with excitation wavelength 485 nm and emission wavelength 525 nm, both with a bandwidth of 15.0. The gain was 75% of the optimal value of the baseline signal (initial time point). Fluorescence measurements were taken every 15 minutes for a total of 120 minutes.

### Immunocytochemistry (ICC)

For monolayer staining, cells were transferred into 8-well glass slides (Millicell EZ Slide, Millipore Sigma #PEZGS0816) precoated with Matrigel at 40,000 cells were plated per well. Cells were fixed with ice cold 4% Paraformaldehyde solution (PFA) (Thermo Scientific #28908) for 30 minutes at room temperature, followed by 3 times washing with PBS. Blocking buffer consisted of 2% goat serum, and 0.25% Triton X-100 in 1X PBS for 1 h at room temperature. All primary antibodies were diluted in blocking buffer according to the manufacturer’s recommended ratio and incubated 16-24 hours 4 °C. After washing 3 times with 1X PBS, the wells were incubated in host species selective secondary antibodies and DAPI nuclear stain for 1 h at room temperature. After washing, glass slides were mounted in Fluoromount-G (ThermoFisher #00-4958-02). Antibodies used include (CD44, ThermoFisher #14-0441-82; GFAP, ThermoFisher #01-670-261; TH, Sigma-Aldrich #T1299; GCase, Bio-Techne #MAB7410; EN1, DSHB #4G11).

For the midbrain organoids, immunolabeling and optical clearing was done according to a previously published protocol (51). Briefly, fixed organoids were treated with washing buffer (OWB) (Triton X-100, SDS, and goat serum in PBS) and mounted in clearing solution (FUnGI, composed of glycerol, H2O, Tris buffer (pH 8.0), EDTA, fructose and urea). The primary and secondary antibodies were both incubated overnight at 4°C while mildly rocking and the organoids were washed 3 times 10 minutes each with OWB between each step.

### Lipidomics

To assess lipidomic changes in reactive astrocytes, 3 separate preparations of cells were treated with or without IL-1α at 10ng/ml (Peprotech #200-01A), TNFα at 30ng/ml (Peprotech #300-01A), and Complement C1q at 400ng/ml (Sigma-Aldrich #C1740) (a.k.a. ITC) to mimic inflammatory conditions. Lipidomic analysis was conducted at the Metabolomics Core Facility at Baylor College of Medicine. Lipid extraction was performed using the modified Bligh and Dyer method. Lipid class content was determined by summing individual lipids that shared the same head group. Enrichment analysis of all normalized lipidomics data was performed via the Lipid Ontology web platform (LION/web) (31) to classify the lipids by their structural and functional properties.

### Transcriptomics

Cells were collected, and RNA was extracted using a Direct-zol RNA Kit (Zymo Research) following the manufacturer’s instructions. The RNA concentration was measured using a benchtop spectrophotometer/fluorometer (Denovix), and reverse transcription was conducted with iScript RT Supermix (Bio-Rad). Quantitative PCR (qPCR) was carried out on the cDNA using SsoAdvanced Universal SYBR Green Supermix (Bio-Rad) on a Fast Real-Time PCR System (7900HT; Applied Biosystems), with GAPDH serving as an internal control for normalization. Changes in gene expression were assessed by calculating fold changes (2−ΔΔCt) relative to control groups, and the primers are listed (**Fig. S4**).

For RNA sequencing, the quality of the purified total RNA was analyzed via microfluidic electrophoresis on a Bioanalyzer 2100 (Agilent). The samples were sent to Novogene Corporation for library preparation, sequencing, and bioinformatics analysis. A threshold of fragments per kilobase of transcript per million mapped reads (FPKM) > 1 was established to determine gene expression. Differential expression analysis was conducted using the DESeq2 R package to identify differentially expressed genes (DEGs). The resulting P-values were adjusted for false discovery rate control using Benjamini and Hochberg’s method (Padj ≤ 0.05). The sequencing data have been deposited in the National Center for Biotechnology Information’s Gene Expression Omnibus under accession number GSE250182.

## ACKNOWLEDGMENTS

Research reported in this publication was supported by the Parkinson’s Progression Markers Initiative (PPMI) (17871), the National Institute of Neurological Disorders and Stroke grant (R01NS12978), and philanthropic funding from Paula and Rusty Walter and Walter Oil & Gas Corp Endowment at Houston Methodist and the Sherman Foundation.

PPMI – a public-private partnership – is funded by the Michael J. Fox Foundation for Parkinson’s Research and funding partners, including 4D Pharma, Abbvie, AcureX, Allergan, Amathus Therapeutics, Aligning Science Across Parkinson’s, AskBio, Avid Radiopharmaceuticals, BIAL, BioArctic, Biogen, Biohaven, BioLegend, BlueRock Therapeutics, Bristol-Myers Squibb, Calico Labs, Capsida Biotherapeutics, Celgene, Cerevel Therapeutics, Coave Therapeutics, DaCapo Brainscience, Denali, Edmond J. Safra Foundation, Eli Lilly, Gain Therapeutics, GE HealthCare, Genentech, GSK, Golub Capital, Handl Therapeutics, Insitro, Jazz Pharmaceuticals, Johnson & Johnson Innovative Medicine, Lundbeck, Merck, Meso Scale Discovery, Mission Therapeutics, Neurocrine Biosciences, Neuron23, Neuropore, Pfizer, Piramal, Prevail Therapeutics, Roche, Sanofi, Servier, Sun Pharma Advanced Research Company, Takeda, Teva, UCB, Vanqua Bio, Verily, Voyager Therapeutics, the Weston Family Foundation and Yumanity Therapeutics. The content is solely the responsibility of the authors and does not necessarily represent the official views of the National Institutes of Health.

## AUTHOR CONTRIBUTIONS

A.T. and R.K. conceptualized the study. A.T., T.N. and R.K. designed the studies. A.T and T.N. performed all experiments with assistance from S.O. and M.M.D.K. A.T., T.N. and R.K. generated figures. A.T., T.N. and R.K. wrote and edited the manuscript. R.K. acquired funding.

## DATA AVAILABILITY

Transcriptomics data were made available in public repositories. Further inquiries can be made to the corresponding author.

Figure S1. (A) Schematic illustrating cas9-mediated insertion/deletion site and sequence of GBA^-/-^. (B) ICC staining of GCase protein. (C) Quantification of ICC stain of CD44 and GFAP in differentiated astrocytes.

Figure S2. (A) Representative images of ICC of TH in midbrain organoids comparing the outcome of the induction of transcription factors ASCL1, LMX1B, and NURR1 (ANL) with doxycycline and the addition of TGFβ and dbcAMP. (B) qPCR analysis of Doxycycline induced genes ANL in addition to DA neurons biomarkers FOXA2, LMX1A, and TH.

Figure S3. (A) RNA sequencing volcano plot of GBA^-/-^ astrocytes compared to GBA^+/+^. (B) RNA sequencing volcano plot of GBA^-/-^ astrocytes with and without treatment with Interleukin 1, TNFα, Complement C1q (ITC). (C) Timeline of midbrain astrocytes differentiation protocol and confirmation with En1 ICC. (D) Principal component analysis of lipidomic profiles demonstrates a distinct lipid composition in GBA^+/N370S^ astrocytes compared to GBA^+/+^.

Figure S4. List of forward and reverse primers used for qPCR analysis.

