## Supplementary figures and images for "Glucocerebrosidase Deficiency Dysregulates Human Astrocyte Lipid Metabolism"

### Figure S1

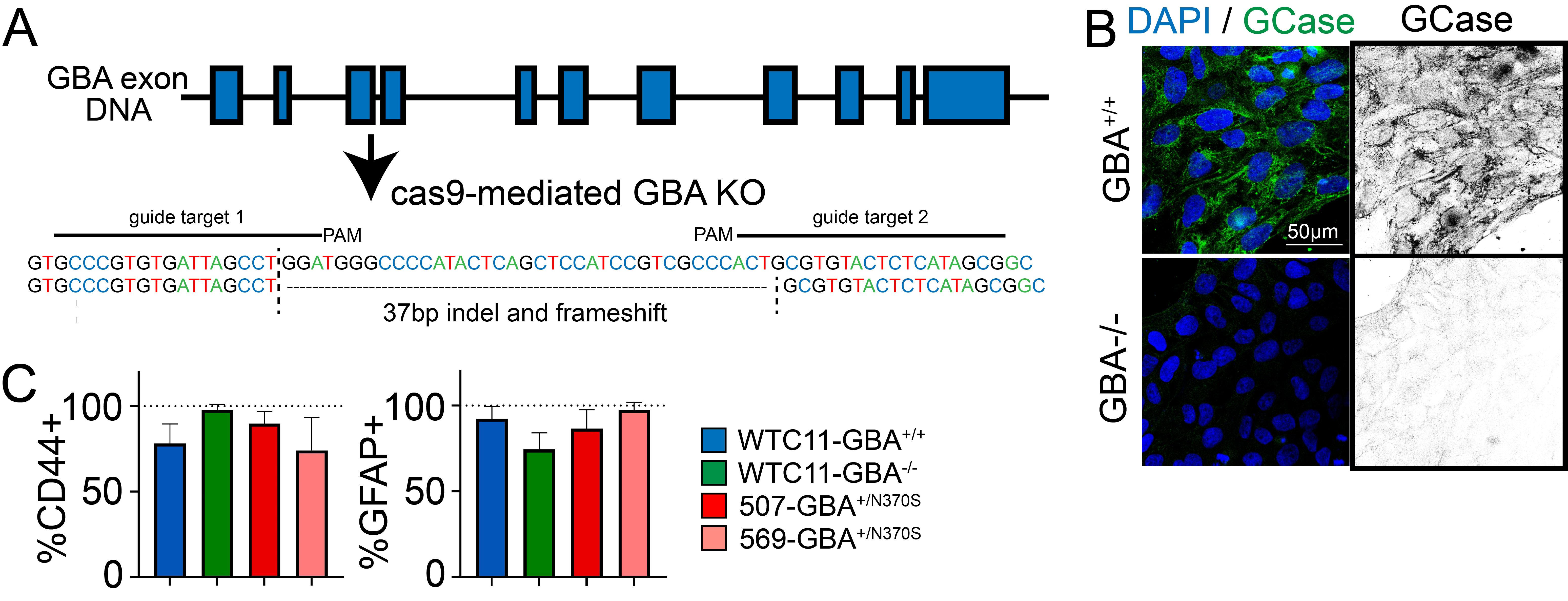

### Figure S2

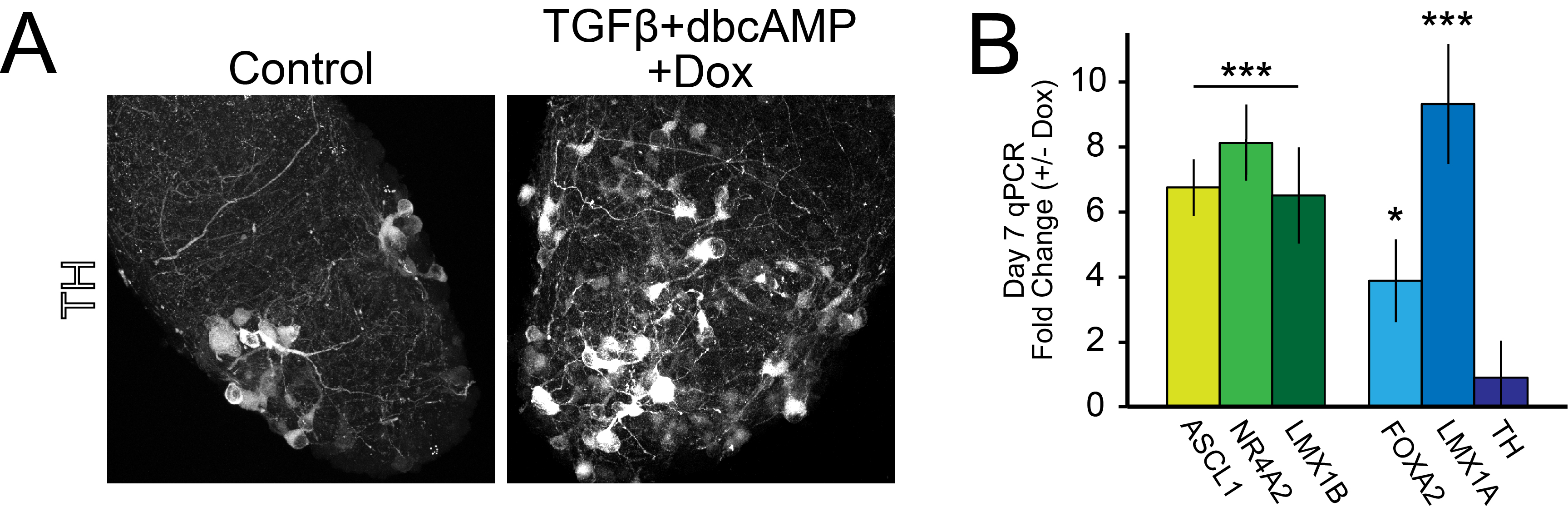

### Figure S3

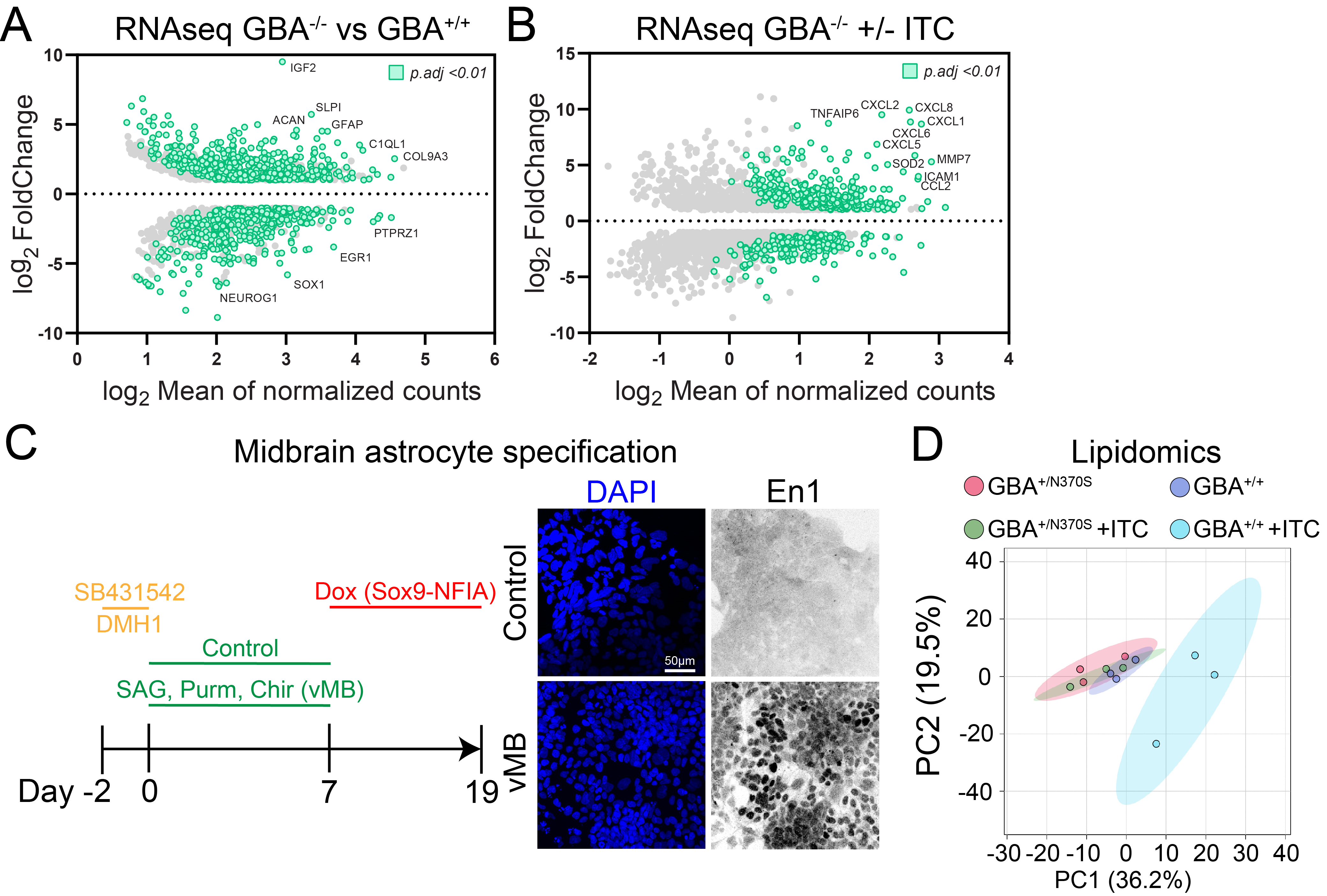

### Figure S4

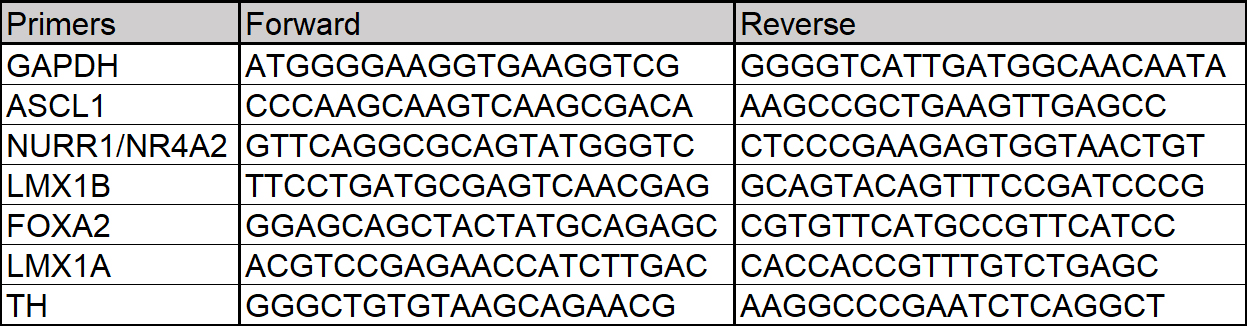
